# Genome-wide libraries for protozoan pathogens drug target screening using yeast surface display

**DOI:** 10.1101/2022.11.17.516844

**Authors:** Rhiannon Heslop, Mengjin Gao, Andressa Brito Lira, Tamara Sternlieb, Mira Loock, Sahil Rao Sanghi, Igor Cestari

## Abstract

The lack of genetic tools to manipulate protozoan pathogens has limited the use of genome-wide approaches to identify drug or vaccine targets and understand these organisms’ biology. We have developed an efficient method to construct genome-wide libraries for yeast surface display (YSD) and developed a YSD fitness screen (YSD-FS) to identify drug targets. We show the robustness of our method by generating genome-wide libraries for *Trypanosoma brucei, Trypanosoma cruzi*, and *Giardia lamblia* parasites. Each library has a diversity of ∼10^5^ to 10^6^ clones, representing ∼6 to 30-fold of the parasite’s genome. Nanopore sequencing confirmed the libraries’ genome coverage with multiple clones for each parasite gene. Western blot and imaging analysis confirmed surface expression of the *G. lamblia* library proteins in yeast. Using the YSD-FS assay, we identified bonafide interactors of metronidazole, a drug used to treat protozoan and bacterial infections. We also found enrichment in nucleotide-binding domain sequences associated with yeast increased fitness to metronidazole, indicating that this drug might target multiple enzymes containing nucleotide-binding domains. The libraries are valuable biological resources for discovering drug or vaccine targets, ligand receptors, protein-protein interactions, and pathogen-host interactions. The library assembly approach can be applied to other organisms or expression systems, and the YSD-FS assay might help identify new drug targets in protozoan pathogens.

## Introduction

Protein-ligand interactions are at the core of every cellular and molecular process in biology. They involve protein interactions with small molecules, nucleic acids, lipids, or other proteins. The discovery of protein-ligand interactions is essential for understanding signaling and developmental processes, gene regulation, pathogen-host interactions, and, notably, developing drugs and vaccines^1^. However, discovering interacting partners is challenging and often relies on screening approaches, including yeast two-hybrid systems, overexpression systems, affinity chromatography coupled to mass spectrometry, or surface display technologies^2-4^.

Yeast surface display (YSD) or phage display combined with high-throughput sequencing is one of the most successful approaches for protein-ligand discovery, mutational scanning, or protein engineering. Examples of their applications include the identification of glucocorticoid receptors^5^, phage receptors^6^, targets of drugs or inhibitors^7^, vaccine targets^8^, T-cell receptors^9^, and antibody targets^10^. YSD entails the expression of exogenous proteins encoded by DNA libraries on the surface of yeast cells. Each yeast cell expresses ∼10^5^ copies of a single protein on its surface, and thus a large yeast population (∼10^8^) can easily represent a complete genomic library. The exogenous proteins are expressed in-frame with a surface anchoring system, e.g., N-terminal fused anchor proteins (SAG1, SED1), the a-agglutinin display system (Aga1p and Aga2p), or the Flo1p display system^11^ (Fig 1A). These anchored proteins are expressed on the exterior of the cell wall and expose the exogenous proteins for ligand interaction. While this approach has proven helpful, challenges in constructing genomic libraries have halted its broad application for routine protein-ligand discovery. Generating a eukaryote’s genome-wide library requires cloning 10^5^ to 10^7^ genome fragments into expression vectors, depending on fragment length and the organism’s genome size. DNA fragments are usually generated from genomic or complementary DNA (cDNA) via enzymatic digestion or random physical fragmentation and then cloned into expression vectors through ligation with restriction digested or T-tailed vectors^12^. Sub-optimal library construction and expression systems can greatly diminish the efficiency of protein-ligand interaction screens. Inefficient library assembly can result in portions of the genome being absent or underrepresented in the library, thus limiting the scope and reliability of downstream screening applications.

**Figure 1.**
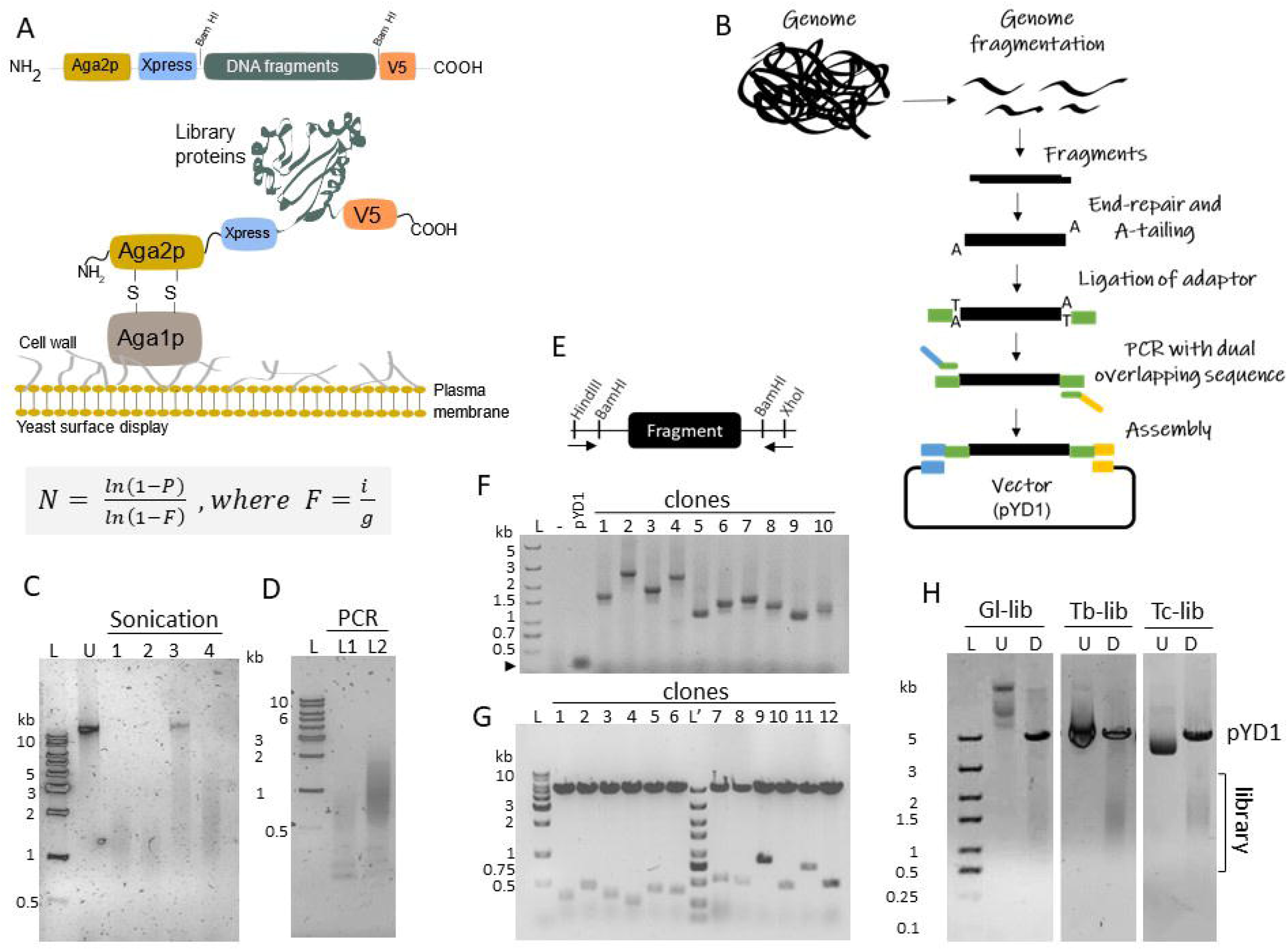
Assembly of genome-wide libraries for YSD. A) Diagram of pYD1 vector (top) showing library fragments cloned within Bam HI sites (by Gibson assembly), and (middle) yeast surface expression using the Aga1p-Aga2p system. Bottom, Clarke and Carbon equation used to estimate the library size, i.e., number of cloned fragments (N) given a probability (P). F is the quotient between insert fragment size (i) and genome size (g) in bp. B) Diagram of the step-by-step genome-wide library preparation. C) *G. lamblia* genomic DNA fragmentation by ultra-focused sonication. DNAs were not sonicated (U) or sonicated at power (W) 25, duty factor (DF) 2%, 200 cycles per burst (cpb), for 45 seconds (s), or 2) 25 W, DF 10%, 100 cbp, 20 s; 3) 10W, DF 2%, 700 cbp, 10s; and 4) 25 W, DF 10%, 200 cbp, 3 s. D) PCR amplification of adapter-ligated library. Library adapter ligation with T4 DNA ligase (L1) or Blunt-TA ligase (L2). E) Diagram of pYD1 vector indicating Bam HI sites flanking cloned fragments and primers (arrows) used for fragment amplification. F) PCR amplification of ten *G. lamblia* library clones (from *E. coli* colonies) to validate Gibson assembly step. The hyphen (-) indicates PCR reactions without colonies. G) DNA restriction analysis of 12 plasmids from *T. cruzi* library clones with Bam HI enzyme to validate Gibson assembly step. Clones are from two libraries with size-selected fragments of ∼300 bp (clones 1-6) and ∼ 700 bp (clones 7-12). H) YSD libraries generated from *G. lamblia* (Gl-Lib), *T. brucei* (Tb-Lib), and *T. cruzi* (Tc-Lib) genomes. Library plasmids were digested with Hind III and Xho I (D). U, undigested library. L or L’ indicate DNA ladders (BioBasic).

The scarcity of genetic tools available to non-model organisms poses significant obstacles to developing genome-wide screenings. This is the case for many protozoan parasites, including kinetoplastids, diplomonads, and apicomplexan, responsible for huge health and economic burden worldwide ^1^. *Trypanosoma cruzi*, the causative agent of Chagas disease (ChD), affects ∼8 million people in South and Central America. ChD has spread to Europe, Australia, and Japan and affects over 300,000 people in the United States ^1^. *Trypanosoma brucei* causes African Trypanosomiasis, also known as Sleeping Sickness, and affects humans and animals with significant health and agricultural impact in Sub-Saharan Africa ^1^. *Giardia lamblia* (also known as *G. intestinalis*) causes intestinal diseases with nearly 200 million yearly cases, especially among children and immunocompromised people in low and middle-income countries ^2^. There are no vaccines against these diseases, and the available drugs have limited efficacy and severe side effects ^1^. While genetic approaches have contributed to understanding *T. brucei* biology and drug-target discovery ^3, 4^, much less has been done for other protozoan pathogens, in which genetic manipulation remains a challenge. Hence, the development of parasite genome-wide libraries for YSD may help advance the knowledge of these organisms’ biology and the development of therapeutics, including identifying drug and vaccine targets.

We have developed an efficient approach to construct genome-wide libraries for YSD. Here, we show the robustness of our method by generating genome-wide libraries for *T. brucei, T. cruzi, and G. lamblia* encoding polypeptides for yeast surface display. The diversity of the libraries ranges from ∼2×10^5^ to 4×10^6^ clones, representing ∼6 to 30-fold of the parasite genome. Nanopore sequencing confirmed the libraries’ genome coverage, and computational analysis predicted the libraries to encode polypeptides in the protein domain range (20-200 amino acids, aa), which is ideal for the identification of ligand binding regions or epitope targeted by antibodies. As a proof-of-concept, we transformed yeast with the *G. lamblia* library and confirmed its polypeptide expression. We developed a YSD fitness screen (YSD-FS) assay for drug target discovery. Using the YSD-FS assay, we identified known interactors of metronidazole, a drug used to treat giardiasis, and new candidate interacting proteins. Notably, the data revealed enrichment in nucleotide-binding domain sequences conferring yeast increased fitness to metronidazole, implying that metronidazole might target multiple proteins containing nucleotide-binding domains. The libraries are valuable biological resources for discovering drug or vaccine targets, ligand receptors, protein-protein interactions, and investigating pathogen-host interactions. The library assembly method can be applied to other organisms or expression systems, and the YSD-FS assay will pave the way to identifying new drug targets in protozoan pathogens.

## Results

### Efficient construction of genome-wide libraries for YSD

To generate genome-wide libraries for YSD, we devised an approach to clone parasite genomic fragments into the pYD1 vector (Fig 1A-B). The method entails genome fragmentation followed by fragments end-repairing and A-tailing for adapter ligation. The adapter-ligated fragments are amplified by PCR with primers that pair with adapter sequences and have ∼20 bp vector overlapping sequences used for fragment cloning by Gibson assembly (Fig 1B). Gibson assembly is a method for joining DNA molecules using a single isothermal reaction and requires a ∼20 bp overlap region between joining molecules ^5^, i.e., fragments and vector. In this approach, a 5’-exonuclease generates 3’-overhangs in the joining molecules, which are then annealed. A DNA polymerase fills gaps between annealed strands, and a DNA ligase seals the nicks in the strands. We initially selected the protozoa *G. lamblia* for genome-wide library construction because it has a haploid genome of 12.1 Mb ^6^, thus smaller than most eukaryotes. This organism’s genome is also primarily organized in arrays of exons with an average gene size of ∼1.5 kb and a coding density of 81.5% of the genome ^6, 7^. The giardia genomic DNA was fragmented in sizes ranging ∼0.5-2 kb by ultra-focused sonication (Fig 1C) to generate an YSD library encoding polypeptides. The genome fragments were end-repaired and A-tailed, followed by ligation of a nanopore forked adapter, which was verified by PCR library amplification (Fig 1B and D). Amplicons from a 16-cycle PCR reaction were used for Gibson assembly reactions followed by bacterial transformation. PCR analysis of randomly selected clones from transformed bacteria confirmed that library clones contained fragments averaging ∼1.5 kb (Fig 1E-F), consistent with the fragmentation, and Sanger sequencing of isolated plasmids confirmed that the sequences originated from the parasite genome. Using Carbon and Clarke equation ^8^ (Fig 1A), we calculated that 36,880 cloned fragments were required to have each *G. lamblia* genome fragment represented in the library with a 99% probability. After library assembly, we obtained ∼2.0×10^5^ clones, which represents 5.3-fold the necessary calculated number of clones with an estimated density of 31 clones per gene (Table 1). Restriction analysis of the constructed library indicated fragments ranging on average ∼1.5 kb (Fig 1H), consistent with fragment size selection.

**Table 1.** Characteristics and diversity of *T. cruzi* (Tc-Lib), *T. brucei* (Tb-Lib), and *G. lamblia* (Gl-Lib) YSD libraries. ^1^ Library sequence mean length calculated from nanopore sequencing data. ^2^ Fold change of the total number of clones obtained compared to the number of clones necessary for one genome coverage at a probability of 99% using Clark and Carbon equation ^33^. ^3^ Library size corrected by percent of coding genome. ^4^Polypeptide length predicted using Libframe tool ^32^.

| **Library Features** | **Tc-Lib** | **Tb-Lib** | **GI-Lib** |
| --- | --- | --- | --- |
| Library sequence mean length<sup>1</sup> | 1057 bp | 644 bp | 860 bp |
| Total number of clones | 4.0x10<sup>6</sup> | 7.2x10<sup>5</sup> | 1.9x10<sup>5</sup> |
| Library genome coverage<sup>2</sup> | 29.6-fold | 6.7-fold | 5.3-fold |
| Calculated number of clones per gene<sup>3</sup> | 270 | 41 | 31 |
| Gene enrichment | 100% | 100% | 100% |
| Predicted protein range (aa)<sup>4</sup> | 13-559 | 13-275 | 13-488 |

To determine the library coverage of the genome, we sequenced the library using Oxford nanopore sequencing. The long-read technology generates sequences covering the complete DNA sequence of the cloned fragments. Nanopore sequencing of the *G. lamblia* library (Gl-Lib) resulted in ∼90% of reads mapped to the *G. lamblia* WB strain reference genome chromosomes and resulted in 27.3x coverage of the genome (Table 2, Fig 2A). Inspection of the parasite chromosomes (chr) showed extensive library representation of the genome with nearly no gap in coverage (Fig 2A and B). The apparent gaps observed in chr 5 relate to the lack of a defined sequence in the reference genome (represented by Ns in the reference genome). About 60% of the library included fragments ranging between 0.4-1.6 kb with an average size of 860 bp (Fig 2C, Table 1). The library read lengths correlated with library fragmentation size, but it also indicated a slight bias towards DNA fragments of small size (Fig 2C, Table 1). Moreover, it also indicated that the library contain fragments to encode predominantly protein motifs or domains ^9^ compared to full-length proteins. The results also revealed that all *G. lamblia* annotated genes were represented in the library (Table 1), and essentially all gene sequences were represented by library fragments (Fig 2D). The data show that the method is efficient for constructing genome-wide libraries.

**Table 2.** Nanopore sequencing metrics of *T. cruzi, T. brucei*, and *G. lamblia* YSD libraries. ^1^Q-score.

| **Library Features** | **Tc-Lib** | **Tb-Lib** | **GI-Lib** |
| --- | --- | --- | --- |
| Total sequenced bases | 8.74 Gb | 3.56 Gb | 1.97 Gb |
| Mean sequencing quality<sup>1</sup> | 12.5 | 11.7 | 12.1 |
| Total number of reads | 8,267,306 | 4,618,344 | 806,545 |
| Sequence genome coverage | 146.9x | 131.9x | 27.3x |

**Figure 2.**
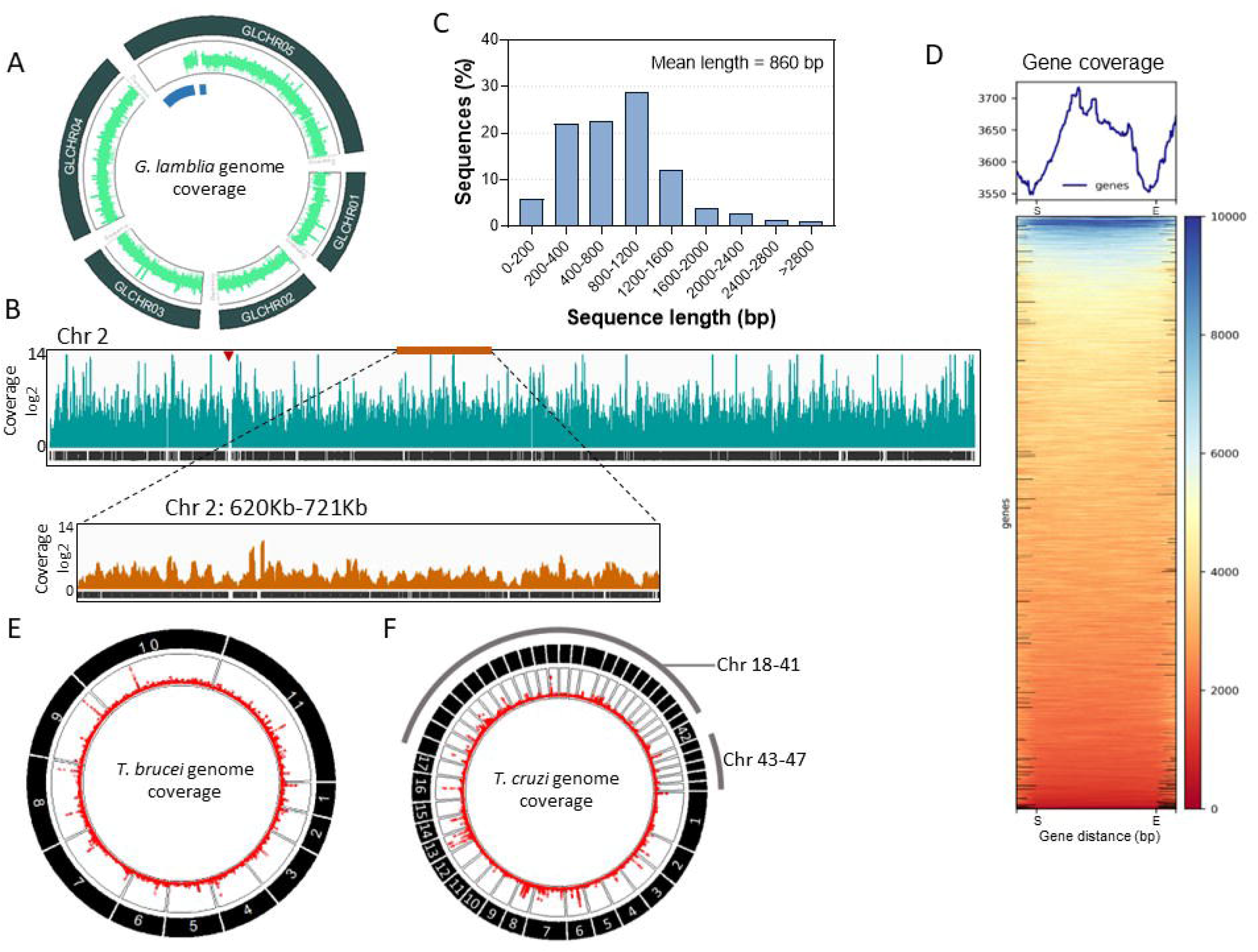
Nanopore sequencing analysis of YSD libraries. A) Circular plot shows the mapping of *G. lamblia* library nanopore sequencing (green) to all five chromosomes (black rectangles). The blue segments indicate the lack of sequence in the reference genome data (hence no mapping). B) Analysis of nanopore read coverage of chromosome 2 (Chr 2). A segment of ∼100 Kb of Chr 2 (orange bar) is highlighted below. Red arrowhead indicates a lack of sequence in the reference genome. C) Distribution of *G. lamblia* genome fragment sequence lengths from nanopore sequencing data. D) Heatmap of all *G. lamblia* genes shows gene read coverage. All gene sequences were resized to 2 kb for read coverage analysis. S, start codon, E, stop codon. A 0.5 Kb upstream (S, gene start) and downstream (E, gene end) is shown. E-F) Circular plot shows the mapping of nanopore sequencing (red lines) of *T. brucei* (E) and *T. cruzi* (F) libraries to the corresponding parasite genome chromosomes (black rectangles).

### Constructing libraries of organisms with large genomes

To further validate our approach, we generated libraries for *T. brucei* and *T. cruzi*, which haploid genomes are 35 Mb and 44 Mb, respectively. These parasite genomes are about 3 to 4 times larger than the *G. lamblia* genome and thus helped test the method’s efficiency and evaluate potential bias associated with genome size. Trypanosome genomes are organized primarily in segments of exons with genes averaging a size of ∼1.5 kb. After library assembly and transformation, we obtained 7.2×10^5^ and 4.0×10^6^ clones for *T. brucei* and *T. cruzi*, respectively (Table 1), representing 6.7 and 29.6-fold the calculated coverage of their genomes at 99% probability, and estimated 41 and 270 clones per gene (Table 1). Restriction enzyme analysis confirmed the library fragment range with an average of ∼1.5 kb (Fig 1G-H). After constructing multiple genome-wide libraries, we found that optimizing the Gibson assembly reaction, which involves defining an optimal ratio between DNA fragments and plasmid vector, is a critical and often necessary step in library construction. After optimization of the Gibson assembly step, we consistently obtained 100% efficiency in library generation using this method. The libraries are hereafter named Tc-Lib and Tb-Lib, respectively, for *T. cruzi* and *T. brucei* libraries.

The nanopore sequencing of Tb-Lib and Tc-Lib resulted in 131.9x and 186.1x genome coverage, respectively (Table 2). Both Tb-Lib and Tc-Lib showed a thorough representation of their respective organism genomes covering all chromosomes (Table 1, Fig2 E-F). The sequencing analysis also revealed an average fragment size of ∼0.7 to 1 kb for the libraries, and every single gene of the parasite’s genome was represented in the libraries (Table 1). There was an apparent enrichment bias for large gene families in all libraries (Fig 2A, E-F). The result is unlikely related to sequence selection during library construction, but it may reflect the poor genome assembly of repetitive regions, especially segments containing large gene families such as mucins or mucin-associated proteins (MASPs) in *T. cruzi*, variant surface glycoproteins (VSGs) in *T. brucei*, and variant surface proteins (VSPs) in *G. lamblia*. The library fragments also covered untranslated regions of the genome. The effective assembly of the three organism’s libraries implies that genome size is not a constraint for library construction by this method, and the data show that the libraries are diverse and comprehensive.

### Library expression in yeast for surface display

We developed a computational tool that predicts the library’s polypeptide sequences potentially expressed on the yeast surface. The libframe tool uses the fastq data from nanopore sequencing. It identifies library sequences in frame with the sequences of interest; here, the Xpress epitope downstream from the Aga2p coding sequence (Fig 1A). Then, it translates in-frame DNA sequences outputting the predicted amino acid sequences and polypeptide length. Analysis of a pool of library sequences indicated polypeptides ranging from 13-488 amino acids (aa), 13-275 aa, and 13-559 aa for the Gl-Lib, Tb-Lib, and Tc-Lib, respectively (Fig 3A, Table 1). Further analysis of the Gl-Lib indicated that 37% of the in-frame library DNA sequences generated predicted polypeptides ranging from 13-39 aa, whereas 67% of sequences encoded proteins ranging from 40-488 aa, which corresponds to about 70,300 and 127,300 clones, respectively, and thus a calculated average of 12 and 21 polypeptides per gene. Notably, over 90% of protein domains range between 20-200 aa with an average of ∼60 aa, whereas motifs and ligand binding sites are 7-8 aa and ∼18 aa, respectively ^9, 10^, implying that the libraries can generate polypeptides encompassing protein functional groups. The library polypeptide range reflects the size distribution of cloned genomic fragments (Fig 2C, Table 1), and early stop codons or frameshifts between library sequences and Aga2p. Early stop codons result from DNA fragments originated from untranslated regions of the genome (18.5% of the *G. lamblia* genome) and coding sequences cloned out of frame, which is common in genomic libraries. Importantly, the large library sizes with hundred thousand to millions of cloned fragments ensure that polypeptides of coding sequences with sufficient length to cover functional protein sequences, motifs, and protein domains are included.

**Figure 3.**
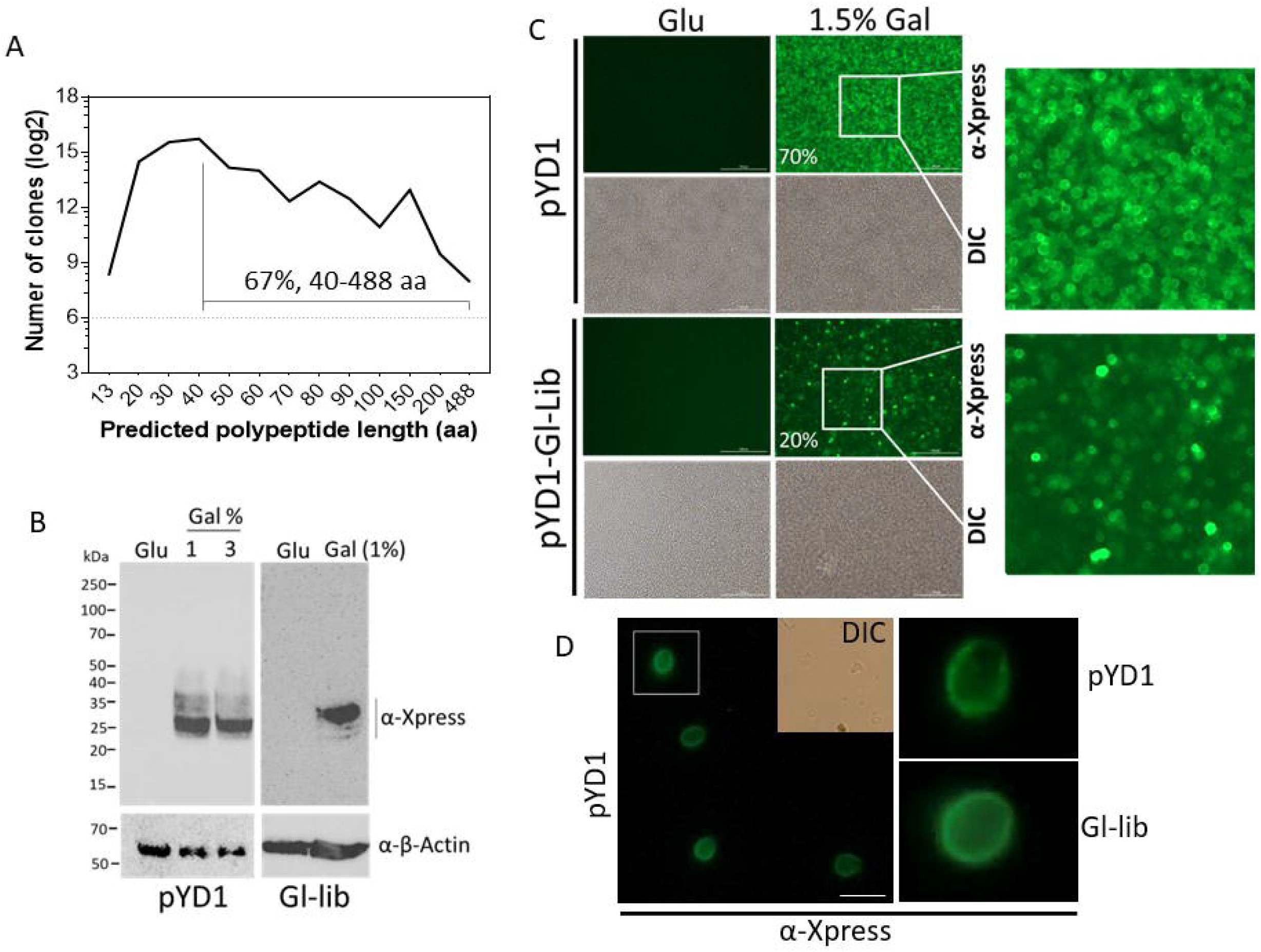
Expression analysis of Gl-Lib YSD. A) Length distribution analysis of Gl-Lib predicted unique polypeptides expressed in-frame with Aga2p using the libframe tool ^32^ (https://github.com/cestari-lab/Libframe-tool). B) Western blot analysis of yeast EBY100 strain transformed with pYD1 (left) or Gl-Lib (right). Proteins were resolved in 12% SDS/PAGE, transferred to nitrocellulose membrane, and blotted with anti-Xpress antibodies. Membranes were stripped and reblotted with anti-β-actin antibodies. Glu, non-induced (2% glucose); Gal, induced with 1-3% galactose in medium containing 2% raffinose. C) Image reader analysis of pYD1 or Gl-Lib transformed yeast. Image analyzed with Cytation 5 image reader in 96-well plates. Bar, 100 µm. Cells were stained with mouse anti-Xpress and AlexaFluor-488-labelled anti-mouse IgG (green). Quantification of high Xpress positive cells by flow cytometry is indicated in %. Glu, 2% glucose; Gal, galactose. D) Fluorescence microscopy analysis of yeast expressing pYD1 or Gl-Lib induced with 1.5% galactose. Cells were treated as described in C). DIC, differential imaging contrast. Bar, 10 µm. Images comparing pYD1 and Gl-Lib from C or D were taken with same light exposure and acquisition time.

We transformed the *Saccharomyces cerevisiae* EBY100 strain with the Gl-Lib or the pYD1 vector to validate library expression. To represent a library diversity of ∼10^6^ clones, high-efficiency yeast transformation is required to obtain at least 10-fold of the library size. Thus, we optimized library transfection in yeast by electroporation and obtained a transformation efficiency of ∼10^7^ cfu/µg of library DNA, which resulted in transformed yeast amounting to ∼230-fold the Gl-Lib diversity. The pYD1 vector has an N-terminal Xpress epitope in frame with Aga2p and expressed fused to the library sequences under control of a promoter induced by galactose (gal) or repressed by glucose (glu) (Fig 1A). Western analysis of gal-induced Gl-Lib confirmed the expression of fragments from ∼25-35 kDa (Fig 3B). The size range correlates with our computational prediction of expressed polypeptides, as most library fragments are predicted to generate 2-7 KDa polypeptides, and the predicted molecular weight of Aga2p plus the Xpress epitope is ∼24 kDa ^11^. We analyzed the expression of the Gl-Lib using anti-Xpress antibodies in a Cytation 5 cell image reader (Fig 3C). The data showed a uniform detection of surface Xpress from pYD1 transformed yeast, and flow cytometry quantification showed that 70% of the yeast uniformly expressed high levels of Xpress epitope. In contrast, the Gl-Lib expressing yeast showed a heterogeneous detection of Xpress (20% strongly positive by flow cytometry), indicating a mixed population of high and low-expressing cells. The low percentage of Xpress positive cells in the Gl-Lib likely reflects interference of the library polypeptides with anti-Xpress antibodies binding to the adjacent Xpress epitope. The data is consistent with other YSD libraries generated using the Aga1p-Aga2p expression system ^12^. Microscopy analysis further confirmed the surface Xpress exposure, indicating a functional yeast surface display system (Fig 3D). The data show that our methodology generates diverse and functional genome-wide YSD libraries.

### Screening for *G. lamblia* proteins that interact with metronidazole

We posited that YSD libraries are useful tools for drug target screenings. The yeast surface expression exposes library proteins to small ligands outside the cell, favouring ligand-protein interaction. The interaction of library proteins with drugs may lead to drug depletion, drug enzymatic inactivation or drug activation into toxic derived molecules, all of which may affect the growth of interacting yeast clones in the population. Hence, we performed a YSD-FS by growing yeast in the presence or absence of metronidazole to identify clones showing altered fitness in the presence of the drug. Gl-Lib expressing yeasts were grown for three days in vehicle only or in the presence of metronidazole, and the grown yeast population was submitted to three rounds of fitness selection in metronidazole. The library plasmid DNAs were recovered from yeasts for fragment DNA sequencing using Oxford nanopore technology. Sequencing analysis of the vehicle-treated YSD showed a 0.8 Pearson correlation with the non-transformed (original) library, implying only a slight deviation in the yeast expressed library from the generated library (Fig 4B, see No screen). However, drug-treated Gl-Lib expressing yeast showed a low correlation with the vehicle-treated library (Pearson correlation of 0.6), implying that drug treatment altered the fitness of the yeast population expressing the Gl-Lib (Fig 4B, MTZ screen). Analysis of metronidazole treated vs non-treated Gl-Lib expressing yeast revealed 270 genes enriched and 26 depleted (Fig 4C, fold-change ≥ 3, *p*-value ≤ 0.01). Of these genes, 108 genes (96 enriched and 12 depleted) had annotations other than hypothetical proteins and included bonafide interactors of metronidazole such as thioredoxin reductase, purine nucleoside phosphorylase, and a putative ortholog of metronidazole target protein 1 ^13^. The screen also identified 13 kinases, primarily from the expanded NEK kinase family; 14 protein 21.1 (Ankyring domain-containing proteins), and genes involved in DNA metabolism, antioxidant metabolism, and membrane transporters (Table 3 and Supplementary Table 1). Importantly, analysis of the enriched library fragments revealed specific gene sequences associated with yeast-increased fitness to metronidazole (Fig 4D). The sequences matched primarily to nucleotide binding regions, e.g., nicotinamide adenine dinucleotide (NAD)-binding domains, P-loop containing nucleotide-binding (NTPase) domain, or ATPase domains, indicating that metronidazole may interact with different proteins containing nucleotide-binding domains, i.e., ATP-binding domains or nucleotide cofactor domains (Fig 4D). Gene ontology (GO) analysis of the annotated genes showed enrichment in heterocyclic compound binding, small molecule binding, purine nucleotide binding, transferase, and protein kinase activities (Fig 4E), confirming a bias towards nucleotide-binding proteins in the genes identified. Analysis of the protein domains encoded by the identified genes revealed that 89% had an annotated nucleotide-binding domain (Fig 4E-F). Hence, the screen identified proteins known to interact with metronidazole, potential drug-binding domains, and new candidate proteins to help identify metronidazole targets and mechanisms of action. The data also shows that YSD-FS is a robust approach for drug-target screening without the chemical labelling or modification of drugs.

**Table 3.**
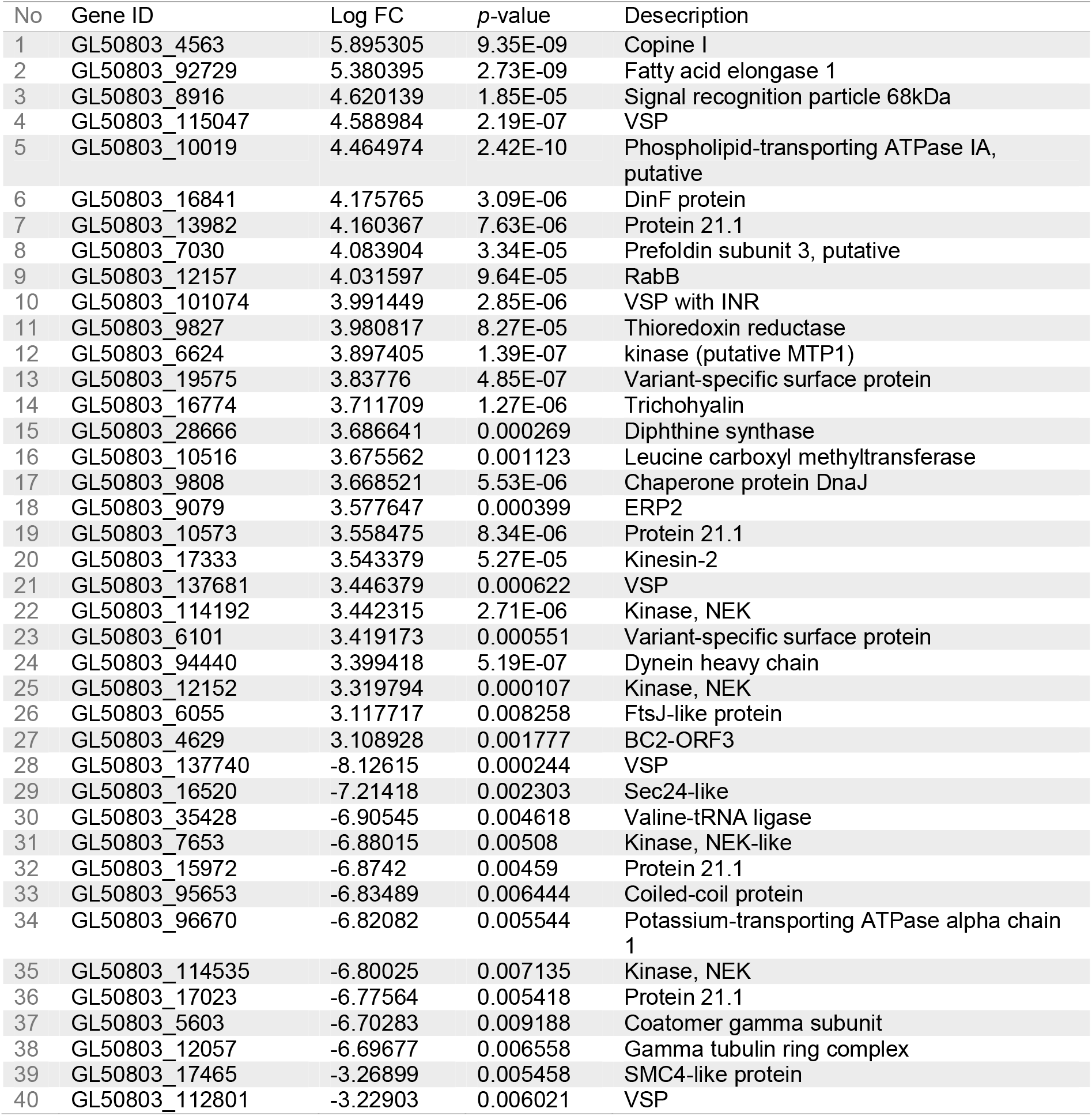
Selection of top *G. lamblia* enriched or depleted genes from metronidazole YSD-FS. Only a selection of 27 enriched and 13 depleted genes with fold-change (FC) ≥ 3 logs and *p-*value ≤ 0.01 is shown. For a complete list, see Supplementary Table 1.

| No | Gene ID | Log FC | $p$-value | Description |
| --- | --- | --- | --- | --- |
| 1 | GL50803_4563 | 5.895305 | 9.35E-09 | Copine I |
| 2 | GL50803_92729 | 5.380395 | 2.73E-09 | Fatty acid elongase 1 |
| 3 | GL50803_8916 | 4.620139 | 1.85E-05 | Signal recognition particle 68kDa |
| 4 | GL50803_115047 | 4.588984 | 2.19E-07 | VSP |
| 5 | GL50803_10019 | 4.464974 | 2.42E-10 | Phospholipid-transporting ATPase IA, putative |
| 6 | GL50803_16841 | 4.175765 | 3.09E-06 | DinF protein |
| 7 | GL50803_13982 | 4.160367 | 7.63E-06 | Protein 21.1 |
| 8 | GL50803_7030 | 4.083904 | 3.34E-05 | Prefoldin subunit 3, putative |
| 9 | GL50803_12157 | 4.031597 | 9.64E-05 | RabB |
| 10 | GL50803_101074 | 3.991449 | 2.85E-06 | VSP with INR |
| 11 | GL50803_9827 | 3.980817 | 8.27E-05 | Thioredoxin reductase |
| 12 | GL50803_6624 | 3.897405 | 1.39E-07 | kinase (putative MTP1) |
| 13 | GL50803_19575 | 3.83776 | 4.85E-07 | Variant-specific surface protein |
| 14 | GL50803_16774 | 3.711709 | 1.27E-06 | Trichohyalin |
| 15 | GL50803_28666 | 3.686641 | 0.000269 | Diphthine synthase |
| 16 | GL50803_10516 | 3.675562 | 0.001123 | Leucine carboxyl methyltransferase |
| 17 | GL50803_9808 | 3.668521 | 5.53E-06 | Chaperone protein DnaJ |
| 18 | GL50803_9079 | 3.577647 | 0.000399 | ERP2 |
| 19 | GL50803_10573 | 3.558475 | 8.34E-06 | Protein 21.1 |
| 20 | GL50803_17333 | 3.543379 | 5.27E-05 | Kinesin-2 |
| 21 | GL50803_137681 | 3.446379 | 0.000622 | VSP |
| 22 | GL50803_114192 | 3.442315 | 2.71E-06 | Kinase, NEK |
| 23 | GL50803_6101 | 3.419173 | 0.000551 | Variant-specific surface protein |
| 24 | GL50803_94440 | 3.399418 | 5.19E-07 | Dynein heavy chain |
| 25 | GL50803_12152 | 3.319794 | 0.000107 | Kinase, NEK |
| 26 | GL50803_6055 | 3.117717 | 0.008258 | FtsJ-like protein |
| 27 | GL50803_4629 | 3.108928 | 0.001777 | BC2-ORF3 |
| 28 | GL50803_137740 | -8.12615 | 0.000244 | VSP |
| 29 | GL50803_16520 | -7.21418 | 0.002303 | Sec24-like |
| 30 | GL50803_35428 | -6.90545 | 0.004618 | Valine-tRNA ligase |
| 31 | GL50803_7653 | -6.88015 | 0.00508 | Kinase, NEK-like |
| 32 | GL50803_15972 | -6.8742 | 0.00459 | Protein 21.1 |
| 33 | GL50803_95653 | -6.83489 | 0.006444 | Coiled-coil protein |
| 34 | GL50803_96670 | -6.82082 | 0.005544 | Potassium-transporting ATPase alpha chain 1 |
| 35 | GL50803_114535 | -6.80025 | 0.007135 | Kinase, NEK |
| 36 | GL50803_17023 | -6.77564 | 0.005418 | Protein 21.1 |
| 37 | GL50803_5603 | -6.70283 | 0.009188 | Coatomer gamma subunit |
| 38 | GL50803_12057 | -6.69677 | 0.006558 | Gamma tubulin ring complex |
| 39 | GL50803_17465 | -3.26899 | 0.005458 | SMC4-like protein |
| 40 | GL50803_112801 | -3.22903 | 0.006021 | VSP |

**Figure 4.**
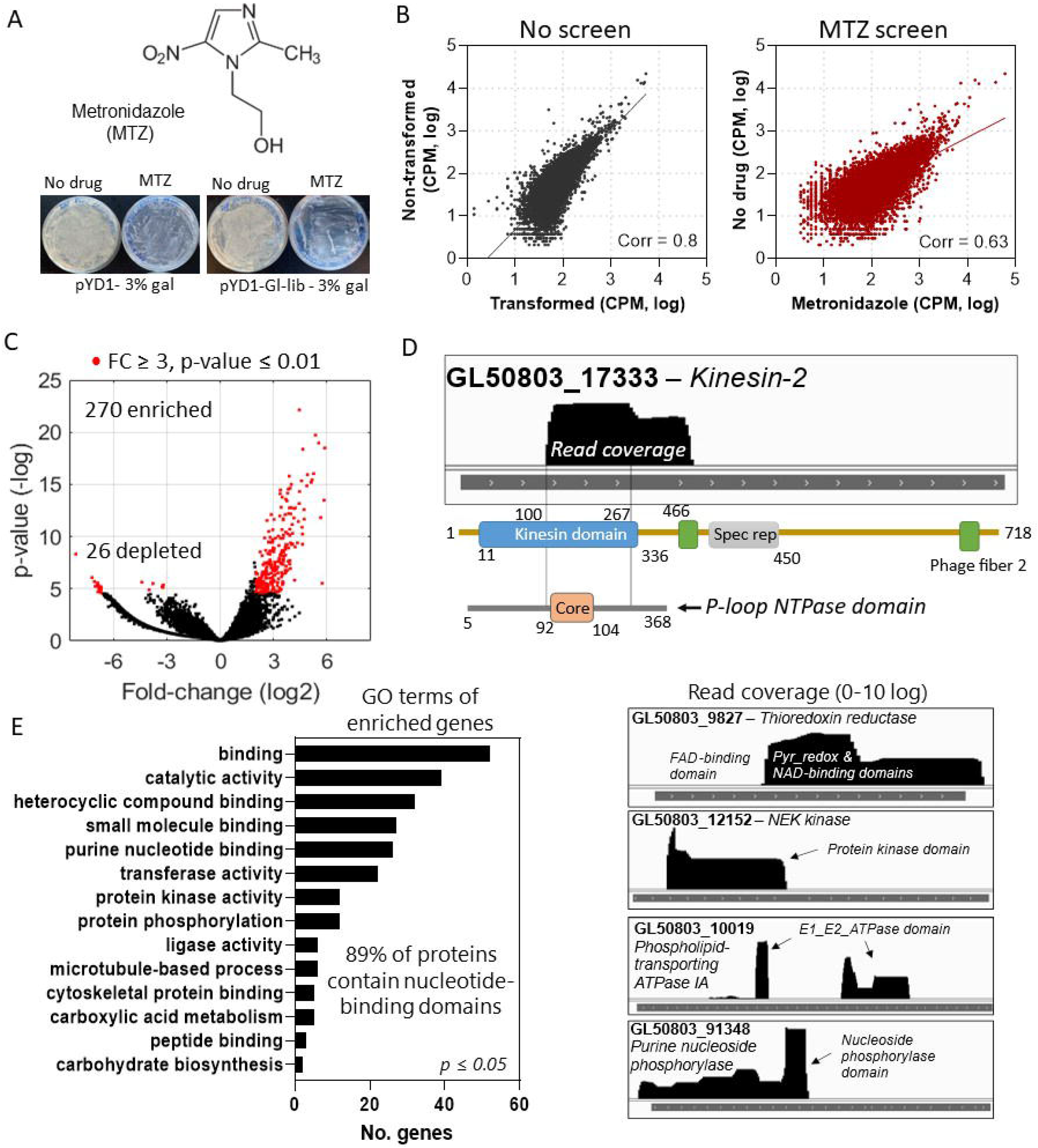
YSD-FS to identify targets of metronidazole. A) Diagram of the drug metronidazole (top). Pictures of Petri dishes with yeast expressing the pYD1 (empty vector) or pYD1-Gl-Lib. Libraries were induced with 3% galactose in the absence or presence of 20 mM of metronidazole. B) Scatter plot of nanopore reads counts per million (CPM) of yeast transformed library with non-transformed library (left, No screen), and yeast expressing Gl-Lib in the absence or presence of metronidazole (right, MTZ screen). Pearson coefficient of correlation (Corr) comparing datasets is shown. C) Volcano plot shows genes enriched and depleted after YSD-FS with metronidazole. FC, fold-change. Statistically significant enriched or depleted hits are indicated in red. D) Read coverage analysis shows the location of library fragments mapped to genes enriched in the metronidazole screen. The coverage was calculated by comparing metronidazole-treated vs non-treated samples. Top, diagram of the kinesin-2 protein shows read coverage mapping to the P-loop NTPase domain. Bottom, read coverage over nucleotide-binding domains in different proteins. E) Go enrichment analysis of the annotated genes enriched in the YSD-FS with metronidazole.

## Discussion

We have developed a robust method to generate genome-wide libraries and demonstrated the method’s efficiency by constructing three genome-wide libraries of protozoan pathogens. We have optimized the libraries’ expression in yeast for surface display and developed an assay combining YSD-FS and high-throughput sequencing to identify potential drug-interacting proteins, demonstrated by using the *G. lamblia* YSD library to screen for potential metronidazole interacting proteins. The screen identified known metronidazole binding proteins, corroborating the assay strength, and new potential metronidazole binding targets. The assay also revealed specific protein domains associated with yeast-altered fitness to metronidazole, indicating potential metronidazole binding sites. The libraries, screening datasets, and methods are valuable resources for identifying drug targets, especially for organisms with limited genetic tools.

Generating eukaryote genomic libraries require cloning between 10^5^ to 10^6^ fragments, which can cause a bottleneck in library construction. To address this problem, we used Gibson assembly to join DNA fragments ^5^, which also facilitates library transfer between different vectors. In our approach, we ligated a forked adapter to the genomic fragments and then used it for fragment amplification with primers containing overlapping sequences to the vector. The ligation of the forked adaptor is straightforward and is routinely used by many laboratories to generate DNA or RNA sequencing libraries. We opted for an Oxford nanopore T/A-based forked adapter because it facilitates the preparation of long-read DNA sequencing of the libraries. In this approach, a single PCR step generates barcoded fragments for DNA sequencing, avoiding long protocols associated with nanopore sequencing library preparation. Still, we anticipate that other adapters, including Illumina or customized adapters, can be used to replace the nanopore adapters during library design. The ∼20 bp overlapping sequence matching the receptor vector (e.g., pYD1) can also be replaced by any vector’s sequence, which provides library transfer flexibility for multiple applications. The timeframe to generate a eukaryote genomic library using this method is about two to three weeks. Other methodologies for library generation include DNA cloning based on ligation of T-tailed vectors or restricted digested DNAs. One limitation of DNA or cDNA restriction enzyme digestion and ligation results from bias in DNA digestion depending on the distribution of restriction enzyme sites in the DNA. At earlier stages of this work, we also attempted generating libraries by DNA ligation using T/A cloning vectors. Although efficient for cloning, custom-made T/A-cloning vectors are not as efficient for large-scale library construction, and they also limit library transfer among different vectors.

We generated three genome-wide libraries with protozoan genomes varying between 12.1 to 44 Mb. There was no bias in library construction related to genome size, and libraries were diverse and complete. Still, a minor tendency toward cloning small fragments was observed, which is common to any cloning approach because small DNA fragments can be overly represented in DNA samples compared to large DNA sequences. The libraries’ high genome coverage and diversity increase the chance that clones containing the desired coding region in frame with Aga2p for yeast surface expression are included. Computational analysis based on nanopore reads predicted thousands of polypeptides in frame with Aga2p. The predicted amino acid sequences included polypeptides in the length range of protein motifs, domains, ^9^ and potential epitopes for protein-ligand binding, including the discovery of antibody targets, protein-domain interaction analysis, and small molecule target screens. We also optimized a protocol for the high-efficiency transformation of yeast and found that the transformed yeast cells preserved the library’s completeness and diversity. We anticipate that the method can help assemble libraries for different applications, including the expression of full-length cDNA sequences, stage-specific transcripts, or gene-specific mutagenesis libraries.

The three libraries generated and validated via nanopore sequencing for *G. lamblia, T. brucei*, and *T. cruzi* are powerful resources for exploring basic and translational biology. Advances in *T. brucei* genetic tools, including RNA interference and genetic screens, have helped identify drug targets and advance our understanding of parasite biology ^3, 14^. However, much less has been done for *T. cruzi* and *G. lamblia*, which have undeveloped genetic tools. RNA interference is present in *G. lamblia* ^15^ but not in *T. cruzi* ^16^, and CRISPR/Cas9 has been used in *T. cruzi* ^17^. However, applying those tools has challenges and limitations, including variability in RNA interference knockdown efficiency, poor parasite transfection efficiency, and lack of regulatable gene expression systems, limiting scaling the approaches for high throughput applications. To date, not a single genetic screen has been performed in *T. cruzi* or *G. lamblia*, which attests to the challenges in the genetic manipulations of these organisms. Hence, harnessing yeast genetics for surface display will help to explore the parasite genomes for drug and vaccine target discovery and advance our knowledge of these organisms’ basic biology.

The YSD technology presents several advantages compared to other protein overexpression systems. Yeast has a cell wall that can limit cell permeability to extracellular ligands and thus prevents potential ligand interactions with intracellular protein targets. Moreover, yeast metabolic enzymes can modify intracellular compounds before interacting with parasite proteins, which are significant limitations of intracellular overexpression screens. However, the surface expression of proteins facilitates ligand-target interactions with little, if any, interference from yeast metabolic enzymes. Furthermore, each yeast cell can express ∼10^5^ copies of a single protein on its surface, which provides multiple ligand interacting sites, and facilitates eukaryotic post-translational modifications of expressed proteins, which are not present in prokaryotic systems such as phage display. The surface display approach can also be used for numerous applications, including ligand-target binding screens, e.g., antibody binding, or phenotypic screens, e.g., drug survival screens.

We performed a proof-of-concept drug screen with yeast expressing the *G. lamblia* library. Although most applications of YSD are ligand-binding, we reasoned that a YSD-FS is more appropriate for drug targets. The YSD-FS does not require drug modifications (e.g., drug labelling with fluorochromes or capturing tags) that can interfere with binding to the unknown target. We used metronidazole because the drug is extensively used to treat *G. lamblia, Trichomonas varginalis*, and bacterial infections ^18^. Moreover, data on a few metronidazole interacting proteins were also available to cross-validate the screen ^13, 19^. The screen identified genes encoding proteins that interact with metronidazole, e.g., thioredoxin reductase and purine nucleoside phosphorylase, thus reproducing described drug targets, and also identified new protein interacting candidates. The annotated candidate interacting proteins’ function was predominantly related to heterocyclic compound binding, with enrichment in proteins with nucleotide-binding domains. The data suggest that metronidazole and perhaps its derived molecules might interact with ATP-binding proteins such as the NEK kinases or enzymes containing nucleotide-binding domains. In agreement with this, metronidazole has been shown to interact with enzymes involved in DNA metabolism ^13, 18, 19^, many of which we identified on the screen. Several Protein 21.1, which are rich in Ankyrin domains and share similarities with the NEK kinases, were identified, likely due to reminiscent kinase/ATP binding domains ^20^. Hence, our data suggest that metronidazole might target multiple proteins sharing a common nucleotide-binding domain. The number of enzymes with nucleotide-binding domains is significant in any organism. The identification of the dozens of enzymes on the screen suggests that amino acid differences within the nucleotide-binding domains affect drug binding. Still, there are some apparent preferences, e.g., the P-loop nucleotide-binding domain. Although further validation of the metronidazole interacting protein candidates will be required, the screen points out directions for this drug targets and mechanisms of action. It might also guide structure-activity relationship experiments to design new target-based compounds. The results also indicate the robustness of the YSD-FS in identifying potential drug interactors at nucleotide resolution.

A potential YSD-FS pitfall is that yeast cells may not be sensitive to the studied drugs. For example, yeast is highly resistant to metronidazole, and the concentrations used in the study differ from those used against *G. lamblia in vitro*. However, the identification of validated metronidazole targets in the screen suggests that the assay tolerates drug ranges different from the concentrations used against the target pathogen. Nevertheless, using mutant yeast strains in multidrug resistance genes may be an alternative in the YSD-FS. Alternating yeast growth conditions from aerobic to anaerobic may also be an alternative to expand the assay application. Notably, the libraries were designed to express polypeptides covering the length of protein domains ^9^ and thus have the advantage of identifying potential drug-binding sites or antibody epitope targets. However, interactions that might require a full-length protein may be missed, as well as proteins or domains not appropriately folded on the yeast surface. The method developed to generate the libraries combined with the YSD-FS and high-throughput sequencing will help expand drug-target discovery and mechanisms of action to organisms in which genetic manipulations are challenging, and tools are not well developed.

## Methods

### Cell culture

*Saccharomyces cerevisiae* strain EBY100 was obtained from the American Type Culture Collection (6). Suspension cultures were grown at 30°C with platform shaking at 225 rpm. Cultures were maintained at 30°C in YPD medium (10 g/L yeast extract; 20g/L peptone; 20 g/L dextrose; pH 6.5) or YPD/agar (YPD with 20 g/L agar). Transformed yeast cells were cultured in a synthetic-defined (SD) medium without tryptophan (SD/-Trp broth, Takara Bio USA Inc.) and induced in SD/-Trp broth with 2% raffinose as an alternative carbon source by the addition of 1 to 3% galactose. *T. brucei* single marker 427 strain bloodstream forms were grown in HMI-9 medium supplemented with 10% inactivated Fetal Bovine Serum (FBS, Life Technologies) ^21^ and 2 µg/mL of G418 (Biobasic) at 37°C with 5% CO_2_. *T. cruzi* Sylvio X10 strain epimastigotes were grown in Liver Infusion Tryptose medium supplemented with 10% inactivated FBS (Life Technologies)^22^ at 27°C. *G. lamblia* WB strain was maintained in modified TY-S-33 medium ^23^ supplemented with 10% inactivated FBS and 2% Diamond’s vitamin solution at 37°C.

### Generation of genome-wide libraries

A detailed protocol for library construction is available ^24^. Briefly, genomic DNAs were obtained from 1.0×10^8^ parasites. Cells were resuspended in 200µl of lysis buffer (10 mM Tris-HCl pH 8, 10 mM EDTA, 1% sodium dodecyl sulfate) and incubated with 20 units of Proteinase K for 30 minutes at 55°C. Afterwards, 20 ul of RNAse A (10 mg/ml) were added, and the samples were incubated at room temperature for 5 minutes. Lysates were mixed vigorously with 1:1 (v:v) Phenol:Chloroform:Isoamyl Alcohol (25:24:1, pH 6.7), followed by centrifugation at 14,000 x g for 10 minutes. The aqueous phase was mixed 1:1 (v:v) with cold isopropanol and 3μl µg of linear poly acrylamide and centrifuged at 14,000 x g for 15 minutes to precipitate DNA. The DNA pellets were washed with 70% cold ethanol and resuspended in 10 mM Tris-HCl pH 8. Next, 2 µg of genomic DNA was fragmented using an ultra-focused sonicator (Covaris M220, Covaris, Inc.) in microTUBE-50 AFA Fiber Screw-Cap tubes. *G. lamblia* genomic DNA was sonicated with peak incident power 25W, duty factor 2%, 200 cycles, for 20 sec at 20 °C. *T. brucei* genomic DNA was sonicated with peak incident power 25W, duty factor 10%, 200 cycles per burst, for 10 sec at 20 °C. *T. cruzi* genomic DNA was sonicated with peak incident power 25W, duty factor 10%, 200 cycles per burst, for 3 sec at 20 °C. DNA fragments of 0.5-3Kb were size selected using Mag-Bind® TotalPure NGS (Omega Bio-Tek) with a beads:sample ratio of 0.5x following the manufacturer’s instructions. DNA fragments were end-repaired and A-tailed using NEBNext Ultra II End Repair/dA-Tailing Module (New England Biolabs Ltd) according to the manufacturer’s instructions, then purified with a 0.5x beads:sample ratio as described above.. The purified DNA was ligated to forked barcode adapters (Oxford Nanopore Technologies) using the NEBNext Quick Ligation Module (New England Biolabs Ltd) for 2h at 20°C. The reaction product was purified as described above. Then, 20 ng of DNA was used for PCR amplification using forward 5’-CGATGACGATAAGGTACCAGGATCCTTTCTGTTGGTGCTGATATTGC-3’ and reverse 5’-TGCAGAATTCCACCACACTGGATTACTTGCCTGTCGCTCTATCTTC-3’ primers. PCR was performed using Taq DNA Polymerase with ThermoPol Buffer (New England Biolabs Ltd) with denaturation at 95° for 3 minutes, followed by 15 cycles of 95°C for 30 sec, 49°C for 30 sec, and 72°C for 3.5 min. PCR products were size selected with 0.5x beads:sample ratio with Mag-Bind® TotalPure NGS (Omega Bio-Tek). Then, 47.04 fmol of fragmented DNAs were used for Gibson assembly with 25.83 fmol BamHI-digested pYD1 vectors using NEBuilder HiFi DNA Assembly Master Mix (New England Biolabs Ltd), and reactions were incubated for 1h at 50°C. One µL of the reaction mix was used to transform 50 µL of chemically competent (∼10^9^ cfu/µg) DH5α *Escherichia coli*. Ampicillin-resistant colonies were scraped from plates, combined, and the plasmid DNAs were isolated using the NucleoBond® Xtra Midi EF plasmid mid-prep kit (Takara Bio USA Inc.) according to the manufacturer’s instructions.

### Nanopore sequencing

Libraries cloned in the pYD1 vector were amplified using forward 5’-TTAAGCTTCTGCAGGCTAGTGGTG-3’ and reverse 5’-CACTGTTGTTATCAGATCAGCGGG-3’ primers with Taq DNA Polymerase and ThermoPol Buffer (New England Biolabs Ltd) for 16 cycles of 95°C for 30 sec, 52°C for 30 sec, and 68°C for 3.5 min. Amplified fragments were purified using 0.85x beads:sample ratio with Mag-Bind® TotalPure NGS (Omega Bio-Tek). Fragments were prepared for Oxford nanopore sequencing using ligation sequencing kit SQK-LSK110 (Oxford Nanopore Technologies) and PCR Barcoding Expansion kit (EXP-PBC001) according to the manufacturer’s instructions, except that quick T4 ligase was used for the BCA ligation instead of the blunt TA enzyme/buffer mix recommended. Fifty fmol of pooled barcoded libraries were sequenced in a MinION using an R9.4.1 flow cell FLO-MIN106 (Oxford Nanopore Technologies) for 24h following the manufacturer’s instructions. A total of 2-8 Gb of DNA were sequenced. All sequences are available at

### Computational analysis

All scripts used in this manuscript are included in the Supporting Information. Nanopore sequencing fast5 data from sequenced libraries were basecalled with Guppy (Oxford Nanopore Technologies), according to the manufacturer’s instructions. Fastq sequences were mapped to *T. cruzi* Sylvio strain reference genome (release 57), or *T. brucei* 427-2018 genome, or *G. lamblia* WB reference genome using minimap2 ^25^. Mapping statistics and format conversions were obtained using Samtools ^26^. Coverage analysis was performed with DeepTools ^27^. Data were visualized using Integrative Genomics Viewer ^28^ and circlize ^29^ following the developer’s instructions. The libframe tool was developed using Python version 3.8 (https://github.com/cestari-lab/Libframe-tool).

### Yeast transfection and galactose induction

EBY100 cells were cultured in Petri dishes containing YPD agar and incubated at 30°C for 72h. In 125 ml of YPD medium, between 5-10 colonies were inoculated to an OD of 0.02 and incubation at 30°C and shaking at 225 rpm overnight until an OD of approximately 1.6. To produce electrocompetent yeast, the cell pellet was harvested by centrifugation (3000 rpm, 4 mins, 4°C) and washed two times in ice cold water, then once in ice cold electroporation buffer (1M sorbitol, 1mM CaCl2). Cells were resuspended in 25 ml conditioning buffer (0.1M LiAc/10mM DTT) for 20 mins at 30°C with shaking at 225 rpm. Cells were harvested by centrifugation and washed once in ice cold water and once in ice cold electroporation buffer. Cells were resuspended in electroporation buffer to a final volume of 1.25 ml and kept on ice until electroporation. For each electroporation, 0.1 ug of library DNA and 5 µl of salmon sperm DNA (ThermoFisher Scientific) (250 µg/ml final concentration) were added to 200 µl of electrocompetent cells and mixed by gently pipetting. The DNA/yeast mixture was transferred to a pre-chilled BioRad GenePulser cuvette (0.2 cm electrode gap) and incubated on ice for 5 mins. Cells were then electroporated at 2.5 kV, 25 μF and 200Ω. Cells were immediately transferred to a 4 ml mixture of 1:1 sorbitol:YPD and allowed to recover for 1 hour at 30°C, shaking at 225 rpm. Cells were collected by centrifugation at 4000 rpm for 5’ at 4 ºC, washed three times in water, and resuspended in 4 ml SD/-trp. 10-fold serial dilutions were prepared; 100 µl of 1:100 and 1:1000 dilutions were plated onto SD/-trp agar and grown at 30°C. The transformation efficiency was calculated from the colony counts after 72 hours. The remaining cells were added to 125 ml SD/-trp per every 200 µl electroporation reaction and the culture was expanded for 24 hours at 30°C with shaking at 225 rpm. Transformed yeasts were then kept frozen in SD/-trp media with 25% glycerol. To induce the library expression of parasite peptides, one mL aliquot of transformed yeast was thawed and resuspended in 30 ml of SD/-trp/2% glucose and grown to an OD600 of 1 at 30 ºC while shaking at 225 rpm. Then, the cells were collected by centrifugation at 4,000 rpm for 5 minutes at 4 ºC, washed three times with cold sterile water and resuspended in SD/-trp/ with 2% raffinose (raf) to an OD600 of 0.5-0.8. Cells were incubated for 4 hours at 30°C with shaking at 225rpm, then collected by centrifugation and resuspended in SD/-trp/2%raf with the addition of 1-3% galactose concentrations for induction of protein expression for 16 hours at 30°C, rotating at 225rpm.

### Western blotting

Protein extracts were prepared by harvesting cells at 5, 000 × g and then resuspending them in 0.1 M NaOH. The cells were incubated at room temperature for 5 min and centrifuged at 5, 000 × g. The pellet was resuspended in 250 mM Tris (pH 6.8), 140 mM sodium dodecyl sulfate (SDS), 30 mM bromophenol blue, 27 μM glycerol, and 0.1 mM dithiothreitol as previously described ^30^. After incubation at 95 °C for 10 min, the samples were centrifuged for 5 min at 5000 g to pellet cell debris. Proteins were resolved in 12% SDS/PAGE and transferred to PVDF membranes as previously described ^31^. Membranes were blocked with 6% non-fat dry milk in PBS 0.05% Tween (PBS-T) for 1h at room temperature (RT). Afterwards, membranes were incubated in mouse anti-Xpress antibodies (Invitrogen) diluted 1:1,000 in blocking buffer (6% non-fat dry milk in PBS-T) for 1h at RT, and then washed 5 times for 8 to 10 minutes at RT. The membranes were incubated in goat anti-mouse IgG-HRP (Santa Cruz) diluted 1:5,000 in blocking buffer for 1h at RT, washed 5 times for 8 to 10 minutes in PBS-T at RT, and then developed with ECL chemiluminescence solution (Life Technologies) in a ChemiDoc Imaging System (Bio-Rad). Membranes were stripped and re-probed, as described above, with rabbit anti-β-actin (1:2000) (ABclonal Technology) and donkey anti-rabbit IgG 1:5000 (Santa Cruz).

### Fluorescence-based imaging and flow analysis

One ml aliquots containing approximately 2.0×10^6^ yeast cells were harvested by centrifugation (4,000 xg, 5 minutes) and washed once in PBS. Cells were then fixed in a 1:1 (vol:vol) mix of PBS and 3.7% paraformaldehyde for 20 minutes at RT on a rotator. Cells were washed three times in FACS buffer (PBS, 1% BSA, 0.1% sodium azide) and resuspended in 1ml FACS buffer. 100 µl of cells were transferred to fresh Eppendorfs, pelleted as above, and resuspended in 100 ul of FACS buffer containing mouse anti-Xpress antibodies diluted 1:500 and incubated at RT for 1 hour or at 4°C overnight with rotation. Cells were then washed three times in FACS buffer and incubated with AlexaFluor 488 conjugated anti-mouse IgG (Invitrogen) in FACS buffer for 1 hour at RT in the dark, with rotation. Cells were washed three times in FACS buffer, then resuspended in 1ml FACS buffer for analysis. Analysis was performed using Attune NxT Flow Cytometer (ThermoFisher Scientific). For Cytation 5 imaging analysis and microscopy, cells were fixed and incubated as described above. Imaging analysis was performed using Cytation 5 imaging reader (BioTek). For microscopy, yeast cells were fixed on cover glass pre-soaked with 1% poly-L-lysine to help cell attachment to the cover glass. Cells were mounted on slides with Fluoromount-G™ Mounting Medium (Thermo Scientific). The slides were analyzed using Nikon E800 Upright fluorescence microscopy (Nikon).

### YSD-FS screen of metronidazole

*Gl-Lib or pYD1* transformed yeast were inoculated into SD/-trp/2%raf/3% gal and incubated at 30°C, 225 rpm for 16h. The cultures were diluted to obtain an OD of 0.2, then 200 ul was inoculated on sterile dish plates (150mm x 25mm) containing SD/-trp/2% raf/3% gal/agar medium and metronidazole (20mg/ml, diluted in dimethyl sulfoxide, DMSO). Cultures were also incubated in plates with drug vehicle only to monitor the effects of DMSO. After a 3-day incubation at 30°C, the colonies from Gl-Lib or pYD1 transformed yeast grown in the presence of metronidazole were scraped from the plates. The cultures were recovered in SD/-trp medium for 16h at 30°C, then washed three times in Milli-Q water and replated as described above in metronidazole-containing plates for an additional three rounds of selection. Afterwards, colonies were scraped from the plates, and their plasmid DNAs were extracted by resuspending the colonies in 400 µL lysis buffer (10mM Tris pH8.0, 0.1M EDTA, 0.5%SDS), 200 μl of acid-washed 0.5 mm glass beads. The mixture was incubated with 20 units of Proteinase K for 30 minutes at 55°C. Then, it was vigorously vortexed for 10 min, with the subsequent addition of another 200µl lysis buffer and vortexed for 10 additional minutes. Afterwards, 20 µl of RNAse A (10mg/ml) were added to the mixture and incubated at RT for 5 minutes. The mixture was boiled at 100 °C for 3 min in a heat block, placed on ice for 1 minute, and then centrifuged for 14,000 × g, 10 min. The supernatant was collected for plasmid DNA extraction using 1:1 (v:v) Phenol:Chloroform:Isoamyl Alcohol (25:24:1) pH 6.7, followed by centrifugation at 14,000 x g for 10 minutes. Cold isopropanol 1:1 (vol:vol) and 3 µl LPA were mixed with the aqueous phase and centrifuged at 14,000 x g for 15 minutes. The precipitated plasmid DNA was washed in 70% cold ethanol and resuspended in 10 mM Tris pH 8.0. The screen was performed in triplicate and samples multiplexed for Oxford nanopore sequencing as described above.

## Data and statistical analysis

Data are shown as means ± standard deviation of the mean from at least three biological replicates. Comparisons among groups were made by a two-tailed t-test using GraphPad Prism. P-values ≤ 0.05 with a confidence interval of 95% were considered statistically significant unless otherwise stated. Graphs were prepared using Prism (GraphPad Software, Inc.), Matlab (Mathworks), or RStudio.

## Data availability

All data have been included within the article and Supplementary Information. Sequencing data were deposited in the Sequence Read Archive (SRA) with BioProject numbers PRJNA851089, PRJNA851424, and PRJNA851903. The Libframe tool and its code is available at https://github.com/cestari-lab/Libframe-tool. The genome-wide libraries are available upon request (Contact: Igor Cestari).

## Supporting information

This article contains supporting information.

## Acknowledgements

We thank Dr. Janet Yee (Department of Biology, Trent University) for providing *Giardia Lamblia* parasites, sharing cell culture protocols, and providing suggestions to the manuscript. We also thank Dr. Steven Rafferty (Department of Chemistry, Trent University) for providing valuable suggestions to the manuscript.

## Funding and additional information

This work was funded by CIHR (CIHR PJT-175222, to IC), NSERC (NSERC RGPIN-2019-05271, to IC), CFI (258389, to IC), and by McGill University (130251, to IC). Andressa Lira received a Mitacs Global-Link fellowship (IT25163). Sahil Rao Sanghi received a fellowship from the Mitacs Research Training Award program (IT19603).

## References

(1) Altamura, F.; Rajesh, R.; Catta-Preta, C. M. C.; Moretti, N. S.; Cestari, I. The current drug discovery landscape for trypanosomiasis and leishmaniasis: Challenges and strategies to identify drug targets. Drug Dev Res 2020. DOI: 10.1002/ddr.21664.

(2) Kirk, M. D.; Pires, S. M.; Black, R. E.; Caipo, M.; Crump, J. A.; Devleesschauwer, B.; Dopfer, D.; Fazil, A.; Fischer-Walker, C. L.; Hald, T.; et al. World Health Organization Estimates of the Global and Regional Disease Burden of 22 Foodborne Bacterial, Protozoal, and Viral Diseases, 2010: A Data Synthesis. PLoS Med 2015, 12 (12), e1001921. DOI: 10.1371/journal.pmed.1001921.

(3) Baker, N.; Alsford, S.; Horn, D. Genome-wide RNAi screens in African trypanosomes identify the nifurtimox activator NTR and the eflornithine transporter AAT6. Mol Biochem Parasitol 2011, 176 (1), 55–57. DOI: 10.1016/j.molbiopara.2010.11.010. Stortz, J. A.; Serafim, T. D.; Alsford, S.; Wilkes, J.; Fernandez-Cortes, F.; Hamilton, G.; Briggs, E.; Lemgruber, L.; Horn, D.; Mottram, J. C.; et al. Genome-wide and protein kinase-focused RNAi screens reveal conserved and novel damage response pathways in Trypanosoma brucei. PLoS Pathog 2017, 13 (7), e1006477. DOI: 10.1371/journal.ppat.1006477.

(4) Collett, C. F.; Kitson, C.; Baker, N.; Steele-Stallard, H. B.; Santrot, M. V.; Hutchinson, S.; Horn, D.; Alsford, S. Chemogenomic Profiling of Antileishmanial Efficacy and Resistance in the Related Kinetoplastid Parasite Trypanosoma brucei. Antimicrob Agents Chemother 2019, 63 (8). DOI: 10.1128/AAC.00795-19.

(5) Gibson, D. G.; Young, L.; Chuang, R. Y.; Venter, J. C.; Hutchison, C. A., 3rd; Smith, H. O. Enzymatic assembly of DNA molecules up to several hundred kilobases. Nat Methods 2009, 6 (5), 343–345. DOI: 10.1038/nmeth.1318.

(6) Xu, F.; Jex, A.; Svard, S. G. A chromosome-scale reference genome for Giardia intestinalis WB. Sci Data 2020, 7 (1), 38. DOI: 10.1038/s41597-020-0377-y.

(7) El-Sayed, N. M.; Myler, P. J.; Blandin, G.; Berriman, M.; Crabtree, J.; Aggarwal, G.; Caler, E.; Renauld, H.; Worthey, E. A.; Hertz-Fowler, C.; et al. Comparative genomics of trypanosomatid parasitic protozoa. Science 2005, 309 (5733), 404–409. DOI: 10.1126/science.1112181.

(8) Clarke, L.; Hitzeman, R.; Carbon, J. Selection of specific clones from colony banks by screening with radioactive antibody. Methods Enzymol 1979, 68, 436–442. DOI: 10.1016/0076-6879(79)68033-3.

(9) Doolittle, R. F. The multiplicity of domains in proteins. Annu Rev Biochem 1995, 64, 287–314. DOI: 10.1146/annurev.bi.64.070195.001443 From NLM Medline. Lin, M. M.; Zewail, A. H. Hydrophobic forces and the length limit of foldable protein domains. Proc Natl Acad Sci U S A 2012, 109 (25), 9851–9856. DOI: 10.1073/pnas.1207382109 From NLM Medline. Tateno, Y.; Ikeo, K.; Imanishi, T.; Watanabe, H.; Endo, T.; Yamaguchi, Y.; Suzuki, Y.; Takahashi, K.; Tsunoyama, K.; Kawai, M.; et al. Evolutionary motif and its biological and structural significance. J Mol Evol 1997, 44 Suppl 1, S38–43. DOI: 10.1007/pl00000056 From NLM Medline.

(10) Khazanov, N. A.; Carlson, H. A. Exploring the composition of protein-ligand binding sites on a large scale. PLoS Comput Biol 2013, 9 (11), e1003321. DOI: 10.1371/journal.pcbi.1003321 From NLM Medline.

(11) Boder, E. T.; Wittrup, K. D. Yeast surface display for screening combinatorial polypeptide libraries. Nat Biotechnol 1997, 15 (6), 553–557. DOI: 10.1038/nbt0697-553 From NLM Medline. Kieke, M. C.; Cho, B. K.; Boder, E. T.; Kranz, D. M.; Wittrup, K. D. Isolation of anti-T cell receptor scFv mutants by yeast surface display. Protein Eng 1997, 10 (11), 1303–1310. DOI: 10.1093/protein/10.11.1303 From NLM Medline.

(12) Bidlingmaier, S.; Liu, B. Construction of yeast surface-displayed cDNA libraries. Methods Mol Biol 2011, 729, 199–210. DOI: 10.1007/978-1-61779-065-2_13.

(13) Leitsch, D.; Kolarich, D.; Wilson, I. B.; Altmann, F.; Duchene, M. Nitroimidazole action in Entamoeba histolytica: a central role for thioredoxin reductase. PLoS Biol 2007, 5 (8), e211. DOI: 10.1371/journal.pbio.0050211 From NLM Medline.

(14) Cestari, I.; Haas, P.; Moretti, N. S.; Schenkman, S.; Stuart, K. Chemogenetic Characterization of Inositol Phosphate Metabolic Pathway Reveals Druggable Enzymes for Targeting Kinetoplastid Parasites. Cell Chem Biol 2016, 23 (5), 608–617. DOI: 10.1016/j.chembiol.2016.03.015.

(15) Marcial-Quino, J.; Gomez-Manzo, S.; Fierro, F.; Rufino-Gonzalez, Y.; Ortega-Cuellar, D.; Sierra-Palacios, E.; Vanoye-Carlo, A.; Gonzalez-Valdez, A.; Torres-Arroyo, A.; Oria-Hernandez, J.; et al. RNAi-Mediated Specific Gene Silencing as a Tool for the Discovery of New Drug Targets in Giardia lamblia; Evaluation Using the NADH Oxidase Gene. Genes (Basel) 2017, 8 (11). DOI: 10.3390/genes8110303 From NLM PubMed-not-MEDLINE.

(16) DaRocha, W. D.; Otsu, K.; Teixeira, S. M.; Donelson, J. E. Tests of cytoplasmic RNA interference (RNAi) and construction of a tetracycline-inducible T7 promoter system in Trypanosoma cruzi. Mol Biochem Parasitol 2004, 133 (2), 175–186. DOI: 10.1016/j.molbiopara.2003.10.005 From NLM Medline.

(17) Lander, N.; Chiurillo, M. A.; Docampo, R. Genome Editing by CRISPR/Cas9 in Trypanosoma cruzi. Methods Mol Biol 2019, 1955, 61–76. DOI: 10.1007/978-1-4939-9148-8_5 From NLM Medline. Soares Medeiros, L. C.; South, L.; Peng, D.; Bustamante, J. M.; Wang, W.; Bunkofske, M.; Perumal, N.; Sanchez-Valdez, F.; Tarleton, R. L. Rapid, Selection-Free, High-Efficiency Genome Editing in Protozoan Parasites Using CRISPR-Cas9 Ribonucleoproteins. mBio 2017, 8 (6). DOI: 10.1128/mBio.01788-17 From NLM Medline.

(18) Leitsch, D. A review on metronidazole: an old warhorse in antimicrobial chemotherapy. Parasitology 2019, 146 (9), 1167–1178. DOI: 10.1017/S0031182017002025 From NLM Medline.

(19) Leitsch, D.; Kolarich, D.; Binder, M.; Stadlmann, J.; Altmann, F.; Duchene, M. Trichomonas vaginalis: metronidazole and other nitroimidazole drugs are reduced by the flavin enzyme thioredoxin reductase and disrupt the cellular redox system. Implications for nitroimidazole toxicity and resistance. Mol Microbiol 2009, 72 (2), 518–536. DOI: 10.1111/j.1365-2958.2009.06675.x From NLM Medline.

(20) Manning, G.; Reiner, D. S.; Lauwaet, T.; Dacre, M.; Smith, A.; Zhai, Y.; Svard, S.; Gillin, F. D. The minimal kinome of Giardia lamblia illuminates early kinase evolution and unique parasite biology. Genome Biol 2011, 12 (7), R66. DOI: 10.1186/gb-2011-12-7-r66 From NLM Medline.

(21) Hirumi, H.; Hirumi, K. Continuous cultivation of Trypanosoma brucei blood stream forms in a medium containing a low concentration of serum protein without feeder cell layers. J Parasitol 1989, 75 (6), 985–989. From NLM Medline.

(22) Cestari, I.; Ramirez, M. I. Inefficient complement system clearance of Trypanosoma cruzi metacyclic trypomastigotes enables resistant strains to invade eukaryotic cells. PLoS One 2010, 5 (3), e9721. DOI: 10.1371/journal.pone.0009721 From NLM Medline.

(23) Keister, D. B. Axenic culture of Giardia lamblia in TYI-S-33 medium supplemented with bile. Trans R Soc Trop Med Hyg 1983, 77 (4), 487–488. DOI: 10.1016/0035-9203(83)90120-7 From NLM Medline.

(24) Sternlieb, T.; Loock, M.; Gao, M.; Cestari, I. Efficient Generation of Genome-wide Libraries for Protein–ligand Screens Using Gibson Assembly. Bio-Protocol (in press) 2022, 12 (22).

(25) Li, H. Minimap2: pairwise alignment for nucleotide sequences. Bioinformatics 2018, 34 (18), 3094–3100. DOI: 10.1093/bioinformatics/bty191.

(26) Danecek, P.; Bonfield, J. K.; Liddle, J.; Marshall, J.; Ohan, V.; Pollard, M. O.; Whitwham, A.; Keane, T.; McCarthy, S. A.; Davies, R. M.; et al. Twelve years of SAMtools and BCFtools. Gigascience 2021, 10 (2). DOI: 10.1093/gigascience/giab008.

(27) Ramirez, F.; Dundar, F.; Diehl, S.; Gruning, B. A.; Manke, T. deepTools: a flexible platform for exploring deep-sequencing data. Nucleic Acids Res 2014, 42 (Web Server issue), W187–191. DOI: 10.1093/nar/gku365 From NLM Medline.

(28) Robinson, J. T.; Thorvaldsdottir, H.; Winckler, W.; Guttman, M.; Lander, E. S.; Getz, G.; Mesirov, J. P. Integrative genomics viewer. Nat Biotechnol 2011, 29 (1), 24–26. DOI: 10.1038/nbt.1754.

(29) Gu, Z.; Gu, L.; Eils, R.; Schlesner, M.; Brors, B. circlize Implements and enhances circular visualization in R. Bioinformatics 2014, 30 (19), 2811–2812. DOI: 10.1093/bioinformatics/btu393.

(30) Andreu, C.; Del Olmo, M. Yeast arming by the Aga2p system: effect of growth conditions in galactose on the efficiency of the display and influence of expressing leucine-containing peptides. Appl Microbiol Biotechnol 2013, 97 (20), 9055–9069. DOI: 10.1007/s00253-013-5086-4 From NLM Medline.

(31) Cestari, I. Identification of Inositol Phosphate or Phosphoinositide Interacting Proteins by Affinity Chromatography Coupled to Western Blot or Mass Spectrometry. J Vis Exp 2019, (149). DOI: 10.3791/59865.

(32) Sternlieb, T.; Loock, M.; Gao, M.; Cestari, I. Efficient Generation of Genome-wide Libraries for Protein–ligand Screens Using Gibson Assembly. Bio-Protocol (in press) 2022, 12.

(33) Clarke, L.; Carbon, J. Selection of specific clones from colony banks by suppression or complementation tests. Methods Enzymol 1979, 68, 396–408. DOI: 10.1016/0076-6879(79)68029-1.

