## Supplementary Information for "Genome-wide libraries for protozoan pathogens drug target screening using yeast surface display"

This file contains the scripts used for computational analysis in this work.

**Scripts for computational data analysis**

The scripts below were submitted to a cluster from Compute Canada (www.computecanada.ca/) using Linux Ubuntu in Windows Subsystem for Linux. A general file name is given for simplicity, e.g., “GenomeOfReference.fasta” for reference genome, or “reads-map_v1.sam” for mapped output file.

1. Decompressing fastq files and mapping the reads to the genome using minimap2.

```
#Before mapping, files in the *.fasta.tar.gz format must be decompressed using the bash command
tar xvzf yourdata.fastq.tar.gz -C pathdirectory

#This script aligns the reads obtained from the ONT sequencing to the genome of reference, creating a sam file.
module load minimap2/2.24

minimap2 -ax map-ont -k11 -m30 -w7 -I4G -t16 -2 GenomeOfReference.fasta \

/~/data/fastq/*.fastq.gz \

>/~/data/analysis/reads-map_v1.sam
```

2. Obtaining mapping statistics and files conversion from .sam to .bam files. Sorting and indexing .bam files.

```
#Samtools commands extract statistics from the read mapping and will create the bam binary files for further processing.

module load samtools

samtools flagstat /~/data/analysis/reads-map_v1.sam > /~/data/analysis/reads-map_v1-flagstat.txt

samtools stat /~/data/analysis/reads-map_v1.sam > /~/data/analysis/reads-map_v1-stat_lib.txt

samtools view -S -b /~/data/analysis/reads-map_v1.sam > /~/data/analysis/reads-map_v1.bam

samtools sort /~/data/analysis/reads-map_v1.bam > /~/data/analysis/reads-map_v1_sorted.bam

samtools index /~/data/analysis/reads-map_v1_sorted.bam > /~/data/analysis/reads-map_v1_sorted.bam.bai
```

3. Coverage analysis of the genome and read counting.

```
#Using the DeepTools module to perform further analysis and generate visualizations.
module load python/3.8.2
#For these steps a virutal environment was created
source ~/ENV/bin/activate

#plotCoverage generates a curve plot depicting the reads per base and the mean coverage of the genome.
plotCoverage -b /~/data/analysis/reads-map_v1_sorted.bam --labels reads-map_v1 --plotFileFormat pdf --outRawCounts Reads-V1_Lib.txt --numberOfProcessors 8

#Load required packages for readcount analysis (package subread). Read count was used to quantify the number of reads per gene.

module load nixpkgs/16.09
module load gcc/7.3.0
module load StdEnv/2020
module load subread/2.0.3

#featureCounts counts mapped reads for genomic features such as genes, exons, promoter, gene bodies, genomic binds and chromosomal locations. The script below will generate the number of read counts for each exon.
featureCounts -LMO -a GenomeOfReference.gtf -o reads-map_v1_counts.txt -F "exon" -g "gene_id" -s 0 -T4 reads-map_v1_sorted.bam
```

4. Processing the data for visualization of library coverage.

#bamCoverage generates a coverage track by normalizing and binning the reads, producing a *.bw file. The file can be analyzed in a genome visualization tool (e.g., integrated genome viewer such as https://igv.org/app/) for visual validation and to check for potential bias in library coverage.

bamCoverage -b reads-map_v1_sorted.bam -o reads-map_v1.bw **--**binSize 20 **--**normalizeUsing RPKM **--**extendReads 1000 **--**outFileFormat bigwig **--**numberOfProcessors 10

#computeMatrix creates a matrix of scores per region that is necessary as intermediate for the visualization of the data as a heatmap.

computeMatrix scale-regions -b 500 -a 500 **-**m 3000 -R GenomeOfReference.gtf -S reads-map_v1.bw **--**skipZeros **--**sortRegions no **--**transcriptID 'exon’ –o matrix_reads-map_v1-sr1.gz

#plotHeatmap generates the heatmap to visualize gene coverage using the matrix file from computeMatrix.

plotHeatmap -m matrix_reads-map_v1-sr1.gz --outFileName reads-map_v1_heatmap.png --colorMap RdBu --whatToShow 'heatmap and colorbar' --zMin -4 --zMax 4

5. Generate graphs for visualization of read coverage using circular plot in R.

```
#Circlize package was used to generate the circular visualization of genome coverage. First, load R.
module load r

#Initiate R
R

#Install required packages
install.packages("circlize")

#Loading necessary libraries
library(dplyr)
library(circlize)

#Reading data from the plotCoverage output file (reads per base)
Lib_Cov.df = read.table('Reads-V1_Lib.txt', sep = "\t")

#Order the table according to read counts
Lib_Cov.ordered<-Lib_Cov.df[order(Lib_Cov.df$V4), ]

#Filter the data frame to have only the reads from complete chromosomes. This is a useful step in the case of partially sequenced or partially assembled genomes of reference.
Lib_Cov.chrom <- dplyr::filter(Lib_Cov.ordered, grepl('CHR', V1)) %>%
#Converting the scale to log2 to improve visualization (optional)
  dplyr::mutate(V4 = log2(V4)) %>%
# Converting  all cells that became –inf during log conversion to NA.
  dplyr::mutate_if(is.numeric, list(~na_if(., -Inf)))

#Generate the circlize graph
#This first row inicializes the circular graph
circos.initializeWithIdeogram(Lib_Cov.chrom, plotType = NULL, circos.par(gap.degree = 8))

#This chunk generates the most outer ring with the chromosomes tracks
circos.track(ylim = c(0, 1), panel.fun = function(x, y) {
  chr = CELL_META$sector.index #Uses the chromosome names as labels
  xlim = CELL_META$xlim
  ylim = CELL_META$ylim
  circos.rect(xlim[1], 0, xlim[2], 1, col = "lightblue") #Changes the color of the chromosome rectangles
  circos.text(mean(xlim), mean(ylim), chr, cex = 1.5, col = "white",
              facing = "bending.inside", niceFacing = TRUE) #Changes the color of the text and the way it is oriented inside the rectangles.
}, track.height = 0.15, bg.border = NA)

#This chunk adds the counts track
circos.genomicTrack(Lib_Cov.chrom,
                    panel.fun = function(region, value, ...) {
                      circos.genomicPoints(region, value, type = "segment", lwd = 2,
                                           col = "magenta", cex = 0.5, ...)
                      circos.yaxis("right", labels.cex = 0.4, col = "grey", labels.col = "darkgrey")
                    })
#This adds a text in the middle of the circle
text(0, 0, "Genome\ncoverage", cex = 1.5)
```

6. Libframe analysis to calculate the predicted peptide lengths and amino acid sequences.

**Note.** The Libframe tool used in this step was developed in python and is used for pYD1. If using a different expression system, the code can be modified to replace the Xpress tag sequence by any sequence that should be in-frame with the cloned fragments. The tool with code assessable is available at https://github.com/cestari-lab/Libframe-tool. To access the code, open the file using a text editor.

```
#The libframe script finds the Xpress tag sequence in the reads and translates everyhting in frame after the end of the tag and until a STOP codon is found.

#Load python and activate enviroment
module load python/3.8.2
source ~/ENV/bin/activate

#Install biopython
pip install biopython

#Decompress the *.fasta.gz files into *.fasta files and concatenate all the multiple FASTA files.
gunzip *.fastq.gz
cat *.fastq > newfilename.fastq

#Execute libframe. It will output a text file with peptides' length and sequence. It takes three commands: 1) path to fastq file; 2) path to output file (.txt); and 3) the minimum length of a peptide. The Xpress tag has 8 aa (24 bases), plus linker (6 bases, 2 aa), and restriction site (Bam HI, 6 bases, 2 aa) resulting in 12 aa sequence DLYDDDDKVPGS. Hence, we recommend the minimum value of 13 for a peptide.

python3 ./libframe.py path/to/fastq/file/.fastq path/to/output/file/.txt 13

#Example of results:
#Protein length is 127 aa and sequence is: DLYDDDDKVPGSTSHAPGRHGGRRIRLELHLDNFKLLPQLCSSVSADGSPAVPLQPVLPGIRQMLHHSVSIKCADARVARFLWPCPYALHCALLSITSEFAAACGSTIWISVVEFCEISSTVAAARV
#Protein length is 16 aa and sequence is: DLYDDDDKVPGSLCWC
#Protein length is 75 aa and sequence is: DLYDDDDKVPGSFLLVLILRRLLGCLTLIRAERHNRPQQGFVRRTAGVFHPSVATKSTCVFHFCISMYGRICGLL
#Protein length is 25 aa and sequence is: DLYDDDDKVPGSFVGADIAGDRAGK
#Protein length is 73 aa and sequence is: DLYDDDDKVPGSFLLVLILRRLHGCLTLCLRVHTLDTKQPEYPLLGGISRAPAHKGAALGGSFSSGCLHAEEV
```

7. Statistical analysis of metronidazole treated vs non treated samples using edgeR.

```
#Circlize package was used to generate the circular visualization of genome coverage. First, load R.
module load r

#Initiate R
R

#Load required libraries
library (limma)
library (edgeR)

#Create DGEList and statistical design
y <- DGEList(counts=data)
group <- factor(c(1,1,1,2,2,2))
design <- model.matrix(~group)
y <- calcNormFactors(y)

#Generate plot MSD for sample comparisons
plotMDS.pdf <-plotMDS(y, col=c(rep("black",2), rep("red",2)))

#Estimate sample dispersion
y <- estimateDisp(y, design)

#Plot Biological correlate of variation
plotBCV(y)

#Generate statistical analysis using generalized linear model
fit <- glmQLFit(y, design)

# Compare groups treated vs non-treated
qlf.2vs1 <- glmQLFTest(fit, coef=2)
```
